# Topology-aware multiscale modeling of viral genomes reveals stability determinants in circoviruses

**DOI:** 10.64898/2026.03.27.714808

**Authors:** Lucianna H. Silva Santos, Simón Poblete, Sergio Pantano

## Abstract

Understanding how viral genomes are organized under extreme spatial confinement remains a fundamental challenge in structural virology. Icosahedral viruses, despite offering high-resolution capsid structures through cryo-electron microscopy and X-ray crystallography, present a major obstacle: their genomes do not obey icosahedral symmetry and are thus averaged out during standard reconstruction procedures, leaving genome topology largely unresolved. Computational modeling offers a complementary avenue, but existing approaches often rely on simplified polymer representations that fail to capture sequence-specific features and the extreme compaction observed in small DNA viruses. Here, we focus on Porcine Circovirus type 2 (PCV2), a member of the *Circoviridae* family and one of the smallest autonomous mammalian viruses, which packages a circular ~1.7 kb single-stranded DNA genome into a ~20 nm T=1 icosahedral capsid at one of the highest DNA packing densities found in nature. We introduce an integrative methodology combining AI-based structural prediction, lattice Monte Carlo simulations, and multiscale molecular dynamics to generate and simulate three-dimensional topological models of the complete PCV2 virion. Our results demonstrate that multiple distinct genome arrangements can produce virions with indistinguishable external morphology, yet differ substantially in their internal stress distributions and predicted particle stability. These findings suggest that PCV2 populations comprise energetically heterogeneous assemblies with implications for infectivity, uncoating, and environmental persistence, while providing a generalizable framework for modeling genome topology in other confined viral systems.

## Introduction

Understanding how nucleic acids are organized under extreme confinement is a fundamental problem in molecular biology and virology. Viral particles provide natural model systems in which entire genomes are compacted to near–physical limits, stabilized, and delivered with high efficiency. Modeling complete viral genomes within their native capsids offers an opportunity to address fundamental questions regarding DNA organization, confinement, and sequence-dependent interactions, while also informing the development of viral vectors, nanoscale delivery systems, and synthetic encapsulation strategies for nucleic acids (1, 2).

Icosahedral viruses are particularly amenable to structural investigation due to the high symmetry of their protein shells. This symmetry has enabled high-resolution determination of proteinaceous capsid structures by X-ray crystallography and cryo-electron microscopy (cryo-EM) of hundreds of viruses, providing detailed insight into protein architecture and assembly principles across a wide range of viral families (3–5).

In contrast, determining viral genome organization remains challenging. Standard reconstruction procedures in both X-ray crystallography and cryo-EM rely on symmetry operations that inherently average out asymmetric structural motifs. Because viral genomes do not necessarily obey icosahedral symmetry, their density is typically smeared or largely eliminated during reconstruction, hampering direct determination of genome coordinates and masking potentially specific capsid–genome interactions (6–8).

Recent methodological advances, including asymmetric cryo-EM reconstruction and focused classification, have begun to enable the visualization of viral genomes in individual particles (9). However, even when technically feasible, single-particle reconstructions may not fully capture the intrinsic heterogeneity of genome organization. Nucleic acids confined within intrinsically disordered protein regions can adopt multiple energetically accessible conformations that differ in topology, yet are consistent with the same global constraints imposed by capsid geometry and charge distribution. Consequently, experimentally resolved single-particle structures may represent only a limited subset of a wide diversity of viable genomic arrangements (10, 11).

Computational modeling provides a complementary framework to explore genome organization beyond what can be directly observed experimentally. Polymer-based models, coarse-grained (CG) simulations, and hybrid approaches have been used to investigate genome confinement, electrostatics, and capsid–genome coupling in a variety of viral systems (2, 12, 13). Nonetheless, many existing models employ simplified representations of the genome or treat it as a uniform polymer (14), limiting their ability to capture sequence-specific features, topological constraints, long-range interactions, and the extreme compaction observed in small DNA viruses. Despite a few examples of explicit modeling of entire viral genomes within experimentally determined capsids (9, 15), the topology of whole viral genomes remains relatively unexplored.

In this work, we focus on a member of the *Circoviridae* family, which represents the smallest known viral family capable of autonomous replication in mammalian cells (16). The two genera within this family, Circovirus and Cyclovirus, include pathogens of veterinary and medical relevance that infect a wide range of vertebrate hosts despite their small genome (17).

Circoviruses encapsidate a circular single-stranded DNA (ssDNA) genome of approximately 1.7-2 kb within an icosahedral capsid of ~20 nm in diameter, resulting in one of the highest DNA packing densities observed in nature (18). Two major open reading frames dominate the genome: ORF1, which encodes the replication-associated proteins Rep and Rep′, and ORF2, which encodes the capsid protein (Cap), the sole structural component of the virion. Notably, except for a short DNA hairpin characteristic of the rolling-circle replication mechanism, the circoviral DNA contains no secondary structure (19). The viral ssDNA is spontaneously encapsulated within the capsid, presumably via capsid–genome interactions dominated by electrostatic interactions. This extreme compaction raises fundamental questions about genome topology, energetic frustration, and the balance between stability and infectivity in such confined systems (20).

Among a large variety of Circoviruses reported, Porcine circovirus type 2 (PCV2) stands out as one of the most intensively studied members within the family, due to its significant economic impact on the global swine industry. Despite widespread vaccination, PCV2 remains an endemic pathogen, exhibiting substantial genetic variability and adaptive capacity (21). Among several genotypes reported, PCV2d is currently the most prevalent worldwide (22). Given the availability of data reported, among which are high-resolution structures, we adopted this genotype as a case study for modeling the genome topology in a virion.

Structurally, the PCV2 capsid adopts a T=1 icosahedral symmetry composed of 60 copies of the Cap protein. High-resolution cryo-electron microscopy and X-ray crystallography studies have revealed a canonical viral jelly-roll fold with extended surface loops that mediate host interactions and antigenicity (**Figure 1A**). The N-terminal segment contains a disordered arginine-rich region that, together with the basic inner capsid surface, forms a positively charged environment that facilitates tight genome association. Specific capsid-DNA interaction sites, known as packaging signals, have been identified for PCV2 (23) and its closely related Beak and Feather Disease Virus (24), featuring 4 and 3 nucleotides bound to homologous sites on the luminal side of the capsid, respectively. Nevertheless, the internal organization of the viral genome and its coupling to other capsid regions remain poorly understood, particularly with respect to three-dimensional topology and energetic heterogeneity (25). Their minimalist architecture imposes strong evolutionary constraints on genome organization, protein-coding capacity, and virion assembly, making circoviruses paradigmatic systems for studying DNA packing at the physical and biological limits of life.

**Figure 1.**
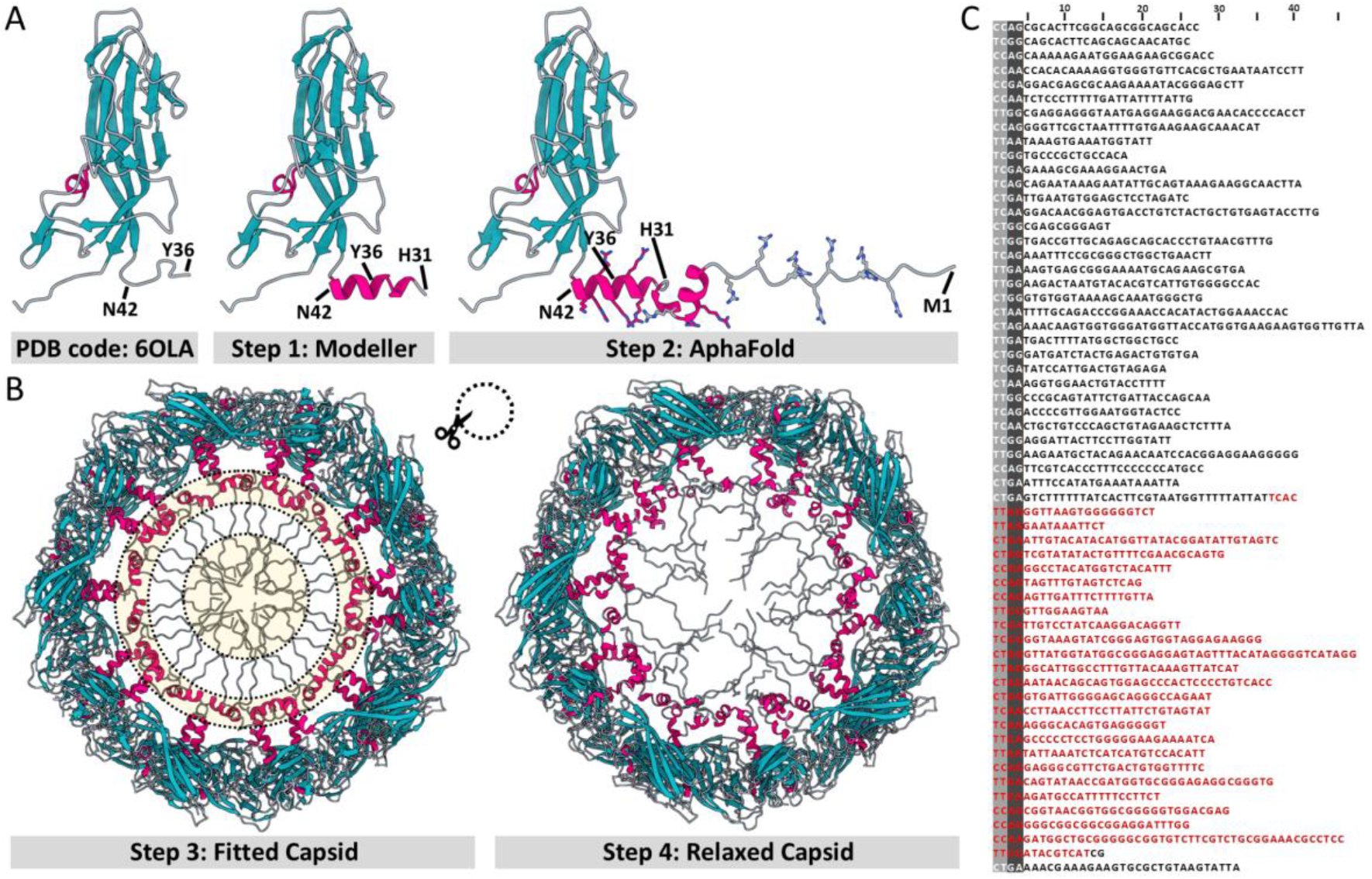
Modeling the PCV2d virion. A) Full-length model of the Cap protein and the PCV2d capsid. The modeling employed a two-step approach, initially integrating a single-turn helix (residues 31 to 42) at the N-terminus. The second step included the remaining missing residues (1 to 30) of the N-terminus. B) The PCV2 capsid is modeled to include the unresolved N-terminal regions. In the fitted capsid (left), internal clashes resulting from the initial superposition of individual Cap proteins are indicated by yellow circles. A relaxed capsid model (right) was obtained using the protocol detailed in the Methods’ section. C) Genome of the PCV2d GeneBank entry KX960929.1. The complete genome sequence is shown, with the putative packaging signals aligned with purines and pyrimidines in gray and black background, respectively. The region encoding the Cap protein, ORF2, is shown in red.

Dissecting the determinants of the intricate network of interactions that leads to such a compact and specific structural arrangement may not only furnish new insights into viral replication mechanisms but also into biotechnological applications for the packaging and delivery of genetic material.

In this study, we introduce a methodology to model the three-dimensional coordinates of the genome inside viral capsids. We combine AI-based structural predictions, lattice Monte Carlo simulations, and multiscale molecular dynamics (MD) simulations to produce and simulate a series of different topological models of the entire PCV2 virion. Our protocol explicitly accounts for the topology of the circular viral genome by integrating experimental data to guide modeling of the genome confined within the capsid. The protocol presented here is summarized in **Supplementary Movie 1, 2** and **3**.

Our results show that multiple genome arrangements can produce virions with indistinguishable overall morphology, yet differ substantially in their internal stress distributions and predicted particle stability. Together, our results provide a unified view of PCV2 virion organization that links genome topology and particle stability and highlights the importance of accounting for physical constraints when interpreting viral genome architecture. These findings suggest that PCV2 particles may exist as a heterogeneous population of structurally comparable but energetically distinct assemblies, with potential implications for infectivity, uncoating, and environmental persistence.

## Methods

Our protocol for modeling entire virions includes modeling the full-length Cap protein, building an energetically relaxed proteinaceous capsid at the fully atomistic level, generating genome topologies, mapping atomistic models to the CG level, and performing MD simulations of the generated models. The entire process is detailed below.

### Modeling the full-length Cap Protein

Due to the absence of residues 1 to 35 at the N-terminus of the PCV2d capsid protein from PDB code 6OLA (23), we employed a two-step approach to model these missing parts. The first step involved modeling residues 31 to 42 of the N-terminus as a single-turn helix, as suggested but not modeled by Khayat et al. (2019) (23). To accomplish this, we utilized comparative modeling with Modeller v10.1 (26, 27), using as a template the bat circovirus structure, PDB code 6RPK (28). The bat circovirus structure contains additional residues at the N-terminus compared to the PCV2 structure, featuring a helix-turn-helix motif that protrudes into the capsid interior.

Having obtained a PCV2 structure with atomic coordinates for residues 31-231, we proceeded to the second step of our approach: modeling the residues before residue 31 using ColabFold v1.5.2 (which uses AF2.3.1) (29). The ColabFold module was used with the PCV2d capsid protein sequence, employing the custom template option to incorporate the comparative modeling model’s atomic coordinates, alongside the multiple sequence alignment option via the MMseqs2-based method (30). In the end, we obtained a PCV2 capsid protein structure with atomic coordinates spanning residues 1-231, as shown in **Figure 1A, right panel**.

### Modeling of PCV2 capsid

The coordinates from our full-length Cap protein were iteratively fitted onto each of the protomers of the molecular envelope of the PCV2d structure, PDB code 6OLA. This resulted in clashes among the 60 N-termini due to their same structural conformation (**Figure 1B**). To address these clashes, the modeled envelope structure was prepared as input for a series of energy minimizations and structural relaxations at 10K temperature, and subsequently at 300K, by employing a Generalized Born (GB) model (31) implemented inside the AMBER22 suite and using the PMEMD module (32). The AMBER FF14SB force field (33) was used to prepare the system topology.

Two energy minimizations were conducted, each comprising up to 1,000 steps. In the first step, strong restraints of 2.4 kcal/mol·Å^2^ were applied to the backbone atoms, whereas in the second step, these restraints were removed. An initial 25 ps run with a time step of 1 fs was used to heat the system to 10K, maintaining all atoms within a restraint of 2.4 kcal/mol·Å^2^. This was followed by two relaxation rounds of 100 ps at a 1 fs time step, reducing the restraint to 0.5 kcal/mol·Å^2^ and then to 0.12 kcal/mol·Å^2^ for the backbone atoms. A final relaxation of 1 ns at a timestep of 2 fs was conducted, applying a restraint of 0.01 kcal/mol·Å^2^ to the backbone atoms. This method effectively addressed proximal side-chain clashes, particularly at the N-termini, without altering the capsid protein’s initial conformation. However, to reorganize the capsid interior and eliminate significant structural clashes, we followed a similar approach at 300K.

At 300K, we used the final frame of the last 10K relaxation to perform two energy minimizations totaling 700 steps. As previously done, the initial minimization imposed a strong restraint of 2.4 kcal/mol·Å^2^ on the backbone atoms, whereas the subsequent minimization was performed without restraints. A heating round lasting 200 ps was employed, maintaining all atoms within a restraint of 2.4 kcal/mol·Å^2^ to gradually heat the system to 300 K. Subsequently, a relaxation round of 200 ps reduced the restriction on the backbone atoms to 1.4 kcal/mol·Å^2^. This was followed by a relaxation round of 200 ps, maintaining a restraint of 1.4 kcal/mol·Å^2^ on non-N-terminal residues and reducing the restraint on N-terminal residues to 0.5 kcal/mol·Å^2^. Another relaxation round ensued, further decreasing the restraint on N-terminal residues to 0.12 kcal/mol·Å^2^ while preserving the 1.4 kcal/mol·Å^2^ restraint on non-N-terminal residues for 500 ps. A concluding relaxation round of 500 ps liberated the N-terminal residues of restrictions while maintaining the 1.4 kcal/mol·Å^2^ restraint on non-N-terminal residues. All relaxation rounds were performed with a time step of 2 fs. The resulting capsid, with a relaxed N-terminus, was used in the subsequent steps for genome insertion (**Figure 1B** and **Supplementary Figure S1**).

### Modeling of the PCV2 genome

Only 60 tetranucleotide Py-Py-Pu-Pu motifs were reported in the 6OLA PDB structure of PCV2d (**Figure 2A**), each of them contacting the capsid’s luminal side. Because of simplicity, the CCGG sequence was assigned to each tetranucleotide (23). However, a simple BLAST search in the GenBank reveals that 60 repetitions of the CCGG motifs are not present in PCV2 genomes. However, the Py-Py-Pu-Pu pattern is found 95 times in the sequence. We therefore clustered these motif occurrences into 60 groups using k-medoids, with sequence distance as the clustering metric, and selected their corresponding centroids as the motifs to be mapped onto the three-dimensional structure. This choice, shown in **Figure 1C**, avoids the selection of closely neighboring fragments and yields a relatively uniform distribution of lengths of the fragments connecting consecutive tetranucleotides. The shortest fragment length found with this procedure was 19 nucleotides, including the packaging signals at the edges.

**Figure 2.**
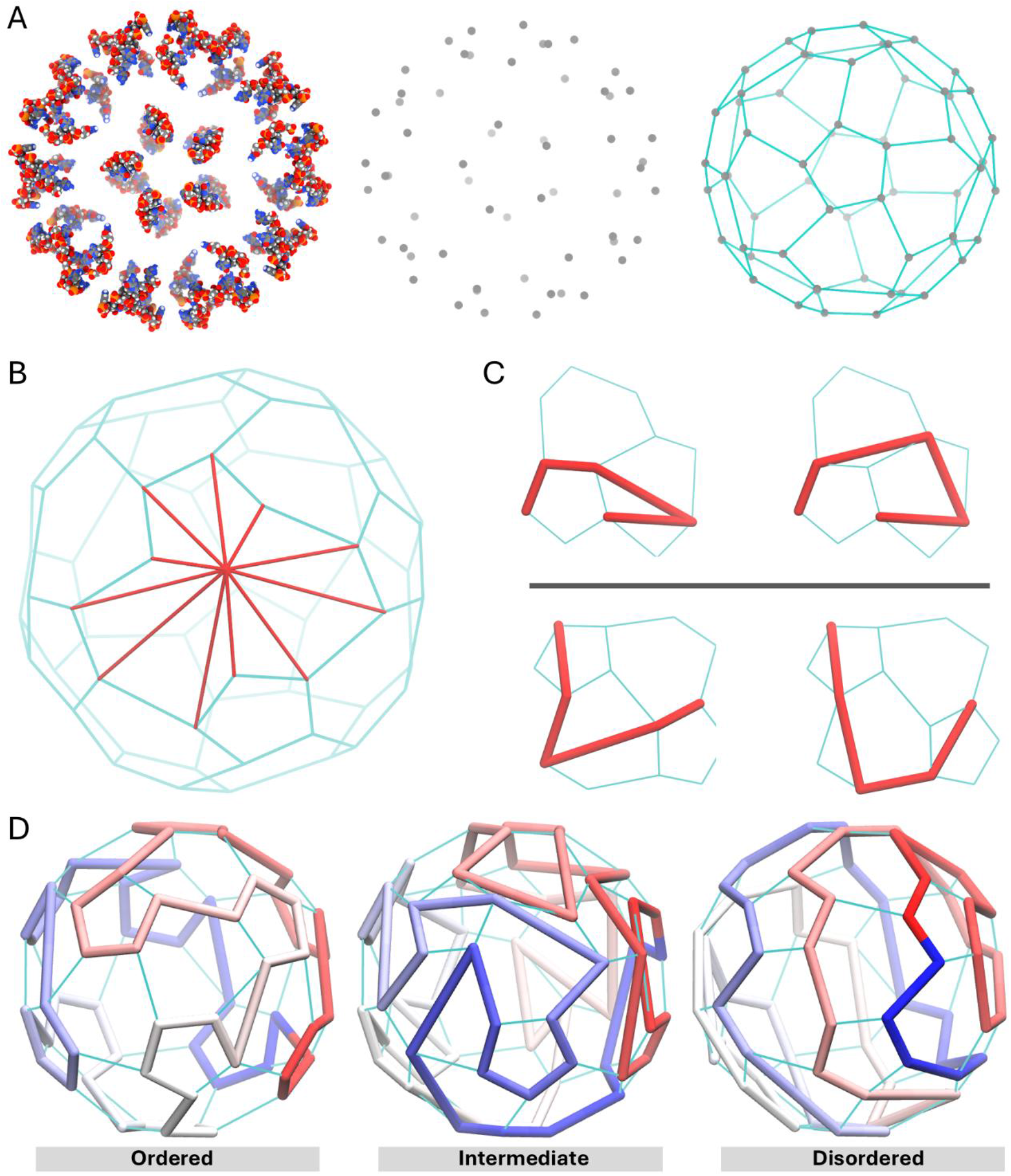
Modeling the PCV2d genome, in its lowest resolution representation. A) The tetranucleotides (packaging signals) determined in the Cryo-EM structure constitute an irregular truncated icosahedron (left) defined by their centers of mass (middle). A lattice-is generated by connecting the centers of mass of the DNA fragments (right). This is illustrated in **Supplementary Movie 1**. B) A lattice point connected with its topological neighbors. C) Monte Carlo trial moves: kink (top) and crankshaft (bottom, see **Supplementary Movie 1**). D) Examples of low-resolution models that illustrate the global topology. The “ordered” model at the left covers all the pentameric units sequentially. The “disordered” model at the right, displays a more extended conformation. At the center, an “intermediate” case generated with the MC procedure.

Nevertheless, the assignment of each tetramer to its corresponding three-dimensional site, as well as the reconstruction details of the connective fragments, is not unique. To address this ambiguity, we implemented a multiscale algorithm that uses a hierarchy of low-resolution genome descriptions to sample and propose a set of complete structures consistent with the available structural data. At the coarsest level, the genome configuration is explored using a lattice-based model (see **Supplementary Movie 1**). The centers of mass of the tetranucleotides constitute an irregular truncated icosahedron, that define possible DNA paths joining its vertices (**Figure 2A**, middle and right panels, respectively). Since all the points are visited once by the path, the resulting configuration corresponds to a Hamiltonian path (34–36). In this representation, the genome can be seen as a polymer constrained to a non-trivial lattice, a perspective that will be exploited to define an efficient protocol for sampling DNA conformations. The low-resolution lattice is then used as a scaffold to construct a nucleotide-level model that introduces sequence specificity. In a final step, pseudoatomistic details can be incorporated to provide a detailed representation of the genome DNA, which is then subjected to a short, localized energy minimization. We describe each step in detail below (see **Supplementary Movie 2** and **3**).

#### Lattice model

The conformations of the genome in the lattice are explored via Monte Carlo (MC) simulations using the Metropolis algorithm: on each integration step, a trial move displaces one or two monomers from their original configuration. Denoting by E_o_ and E_n_ the energies of the old and new configurations, respectively, the trial move is accepted with probability

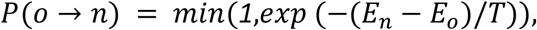

where *T* is the temperature. The trial moves are based on the kink and crankshaft moves used in lattice polymer simulations (37). In the present case, the topology differs substantially from the usual square or triangular lattices used in polymer simulations. So, we have adapted the trial moves accordingly. We begin by defining the connectivity between the lattice sites. Two points are topologically connected if their distance is lower than 5.7 nm, which links the points along the edges and diagonals of the hexagonal and pentameric faces, as shown in **Figure 2B**. The kink moves involve the motion of a single monomer, while the crankshaft moves simultaneously two connected monomers to neighboring sites. In both cases, the trial positions are chosen to maintain the chain’s connectivity, as illustrated in **Figure 2C**.

The energy of a given conformation penalizes the occupation of a lattice site by more than one monomer, the crossings of segments along different diagonals of the same hexagonal face, and small angles formed by three consecutive neighbors. Denoting by *N* the number of monomers, *i, j* the indexes of monomers (ranging from 1 to *N*), and *S*(*i*) and *S*(*j*) the indexes of their lattice sites, the energy *E* of a genome conformation is given by

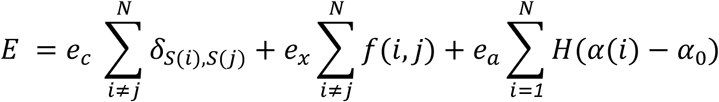

defined in lattice energy units. Where *N* = *60, δ*_*ij*_ is the Kronecker delta, (*i, j*) is a function which is 1 if the segments joining sites *S*(*i*) and *S*(*i* + *1*) and the segment joining monomers *S*(*j*) and *S*(*j* + *1*) are both diagonals of the same hexagonal face and zero otherwise, *α*(*i*) is the angle between the three-dimensional coordinates of sites *S*(*i*), *S*(*i* + *1*), *H*(*x*) is the Heaviside function and *α*_*0*_ = *35*^*°*^. *e*_*c*_, *e*_*x*_, and *e*_*a*_ are parameters set to 10, 50, and 50 lattice energy units, respectively. To ensure the periodicity of the circular genome, the condition *S*(*N* + *1*) = *S*(*1*) and *S*(*N* + *2*) = *S*(*2*) is imposed.

A simulated annealing procedure (38) is applied to 100 replicas, starting from a collapsed state in which all monomers are located at the same lattice point. The annealing procedure consisted of sequential short simulations of 10000 MC steps, in which a trial move is applied to a randomly selected monomer. The initial temperature is 5 lattice energy units, and it is multiplied by a factor *λ* = *0*.*5* if the energy of a short simulation is larger than the energy of the last step of the previous simulation.

#### Nucleotide model

The introduction of the nucleotide-level details is performed independently for each pair of consecutive monomers along the chain of the lattice representation. The correspondence with the lattice representation is established by minimizing the discrepancy between the length of the sequence of the fragments and the three-dimensional distance of their monomers in the lattice. This is performed by normalizing the sequence length of the fragments by their average sequence length, and the three-dimensional length of the segments of the Hamiltonian path by their average over the whole path. For each of the 60 mapping possibilities, the sum of the squared differences of the normalized fragment lengths and normalized segment distances is calculated, and the correspondence with the lowest value of this difference is chosen.

Each fragment is flanked by two packaging signals and mapped to a residue-level CG representation. Because of technical convenience, we used the SPQR model (39, 40). Although SPQR was originally designed for RNA simulations, ssDNA can be readily modeled by imposing the C2’-endo pucker and including only excluded-volume interactions and backbone connectivity. In addition, we apply ERMSD restraints (41–43) (cutoff of 100, coupling constant of 50 *ϵ*_*s*_, denoting by *ϵ*_*s*_ a SPQR energy unit, temperature of 9*ϵ*_*s*_) to the packaging signals from the PDB structure, positioned at the lattice positions. A short simulation of 1000000 MC steps places the packaging signals into the right conformation, and another simulation of the same length minimizes the radius of gyration. Once the 60 segments are arranged, the full genome is assembled. We use energy minimization to remove clashes, freezing the entire structure except for the segments that contain at least one clash. Finally, we run another simulation of 100000 MC sweeps, keeping the packaging signals fixed and minimizing the radius of gyration through harmonic restraints towards an equilibrium value. This compression step exhibited differences in some cases in controlling the degree of compaction, as explained below.

This procedure produces genome conformations at residue-level detail according to the lattice topology derived from the CryoEM data and without clashes.

#### All-atom

An atomistic template of each nucleotide is mounted on top of each SPQR nucleobase. The bond and angle energies are minimized towards the equilibrium values defined in the AMBER force field (44), splitting the genome into strands of 10 nucleotides in order to manually parallelize the energy minimization and significantly reduce the calculation time. The whole procedure can be extended to other geometries if the connectivity map is provided. Other restraints or genome topologies can also be easily incorporated. Arbitrary lattice conformations can also be generated manually or by easily adding energetic terms that favor a particular geometry if experimental information is not available.

To assess the influence of the genome topology on the stability of the virus, we generated three different types of topologies by defining Hamiltonian paths that walk through the lattice visiting packaging signals in different manners. Our classification pays special attention to the way in which the pentagons formed by the five proteins contiguous to a 5’fold symmetry axis are visited by the ssDNA. In “Ordered” genomes, the ssDNA filament visits consecutively each of these 12 pentagons, displaying a locally structured path. In “Disordered” genomes, the ssDNA filament visits all the packaging signals without completing any pentamer consecutively. Therefore, the genomes of this kind exhibit a more extended topology, favoring the connection of distant packaging signals instead of the closest neighbors. Intermediately ordered genomes alternate between both arrangements, having 6 consecutive pentamers visited (I1) and only one, but not along the edges (I2). Examples of these conformations are illustrated in **Figure 2D**.

### Coarse-grained molecular dynamics simulations

Coarse-grained simulations were conducted employing the SIRAH 2.0 (Southamerican Initiative for a Rapid and Accurate Hamiltonian) force field (45), which features a thorough parametrization for DNA-protein interactions (46). MD simulations were performed using the AMBER engine as implemented in AMBER22 (32). The modeled PCV2d virion (capsid and genome) was transformed from a completely atomistic structure to a SIRAH CG representation using SIRAH Tools (47). We employed PACKMOL (48) to incorporate SIRAH’s CG water (49) (named WatFour or WT4) to hydrate the virion’s exterior, with a 3.5 nm radius layer encircling the capsid. Furthermore, to attain an ionic strength of 0.15M, sodium and chloride CG ions were incorporated into this outer layer of WT4. Given that the overall system possesses a net charge of +33 (+1800 capsid and −1767 DNA), we distributed the charge by adding supplementary CG ions to the exterior of the virion to neutralize the system, according to Machado et al. (50), thereby preventing the introduction of ions near the genome. The system derived by PACKMOL was situated within a truncated octahedral computational box via LeaP (51), extending an extra 1.7 nm in all directions of the WT4 outer layer. Consequently, this last step incorporates WT4 molecules into the virion, enhancing hydration near the DNA and yielding a variable amount of WT4 depending on the genomic organization.

Energy minimization and relaxation procedures were conducted following a protocol analogous to that outlined in prior studies (52, 53). Three energy minimizations (EM1, EM2, and EM3) were conducted for all systems for a maximum of 1,000 steps. In EM1, strong positional restraints of 2.4 kcal/mol·Å^2^ were imposed on the capsid proteins and genome. In EM2, the restrictions were applied solely to the backbone beads corresponding to the alpha carbon (GC), nitrogen (GN), and carbonyl oxygen (GO) of the capsid proteins, as well as the backbone phosphate (PX) and backbone C5′ (C5X) of the genome nucleotides. Due to the intricacy and internal arrangement of the systems (N-terminus, genome, and solvent molecules), EM2 and EM3 were repeated to achieve lower energies. The coordinates from the final minimization round were subsequently employed in the initial relaxation round (EQ1) within the NPT ensemble at 300 K, using the middle thermostat scheme implementation with Langevin dynamics (54) and a friction coefficient of 50 ps^-1^. The pressure was isotropically regulated at 1 bar via the Berendsen barostat (55). During this round, the solvent is equilibrated with positional restraints of 2.4 kcal/mol·Å^2^ imposed on both protein and DNA beads. Two successive rounds of side chain relaxation are conducted (EQ2 and EQ3), maintaining the same temperature and pressure controls. Throughout these rounds, the positional restraints were reduced to 0.24 kcal/mol·Å^2^. In the first round, beads of the protein backbone (GN, GO, and GC) and DNA (PX and C5X) were subjected to restraints, whereas in the second round, only the GN and GO beads were constrained. Three additional rounds of side chain equilibration were conducted (EQ4, EQ5, and EQ6), maintaining GN and GO restraints of 2.4 kcal/mol·Å^2^ to reduce the friction coefficient from 50 ps^−1^ to 5 ps^−1^. Before the production phase, an unrestrained 10 ns molecular dynamics relaxation (EQ7) was performed in the NPT ensemble. Again, due to the complexity of the systems’ internal arrangement, reducing the friction coefficient required additional rounds of simulation. However, at least seven rounds were carried out to relax all systems.

All systems underwent unrestricted MD simulation production in the NPT ensemble, maintaining the same temperature and pressure parameters as the preceding relaxation round. All relaxation and production simulation rounds used a time step of 20 fs and a cutoff of 1.2 nm for nonbonded interactions, with snapshots captured every 100 ps. The Particle Mesh Ewald (PME) method (56) for long-range electrostatics was employed and calculated at every integration step. The PME neighbor list was refreshed whenever a bead moved by more than half of a nonbonded “skin” of 0.2 nm. The initial production simulation was run for 2 µs at 300K (27 °C) to establish a stable system. This round was followed by a gradual increase in temperature in five increments of 200 ns each to attain 350K (77°C). The heating simulations were performed to understand the mechanism underlying protein-DNA interaction inside the capsid.

### Structural and Trajectory Analyses

SIRAH Tools GUI package (57) was used to calculate root-mean-square deviations (RMSD), inter-residue (protein-protein) and intermolecular (protein-DNA) contacts, and interbead distances. All plots were created using R (58) and Rstudio (59).

RMSD was calculated using the GC beads of the capsid proteins. Inter-residue and intermolecular contacts were applied to evaluate alterations throughout the simulation at the axes of symmetry and between the genome and capsid, respectively. CG beads from the residues constituting the 2-fold, 3-fold, and 5-fold axes were selected within a radius of 1.5 nm to calculate the contacts using a distance cutoff of 0.8 nm. In contrast, all beads were used to assess protein-DNA interactions, with a cutoff distance of 0.6 nm. The protein-DNA contacts were averaged across the 60 copies of the capsid protein to calculate the percentage of nucleotide-protein residue contacts. This information was also used to establish which nucleotides formed contacts with a protein residue.

The arithmetic average of interbead distances was employed to determine the capsid diameter and the distortion of the 5-fold axes pore. The diameter of the capsid was determined by measuring the distance between K58 residues of capsid proteins situated on opposing sides of the capsid.

Total energy of the systems and protein-DNA energy were calculated using the energy command of CPPTRAJ (51, 60).

## Results

### Hamiltonian path-defined genome topologies influence capsid topology

As briefly explained in the Methods section, integrating information about icosahedral symmetry, the positions of packaging signals, and the circular nature of the genome yields valuable insights into possible topological arrangements. However, our modeling procedure also revealed that a wide conformational diversity is still attainable. Because of simplicity, we restricted our analysis to three groups, namely, i) Ordered: the genome visits each of the DNA-binding sites in a locally ordered fashion within a single Cap pentamer to then pass to the next one; ii) Intermediately ordered: the ordered fashion is respected at least for one pentamer, and iii) packaging signals are visited in a disordered fashion, eventually jumping from one pentamer to a non-contiguous one (**Figure 2D**). In all cases and all the models, all packaging signals are bound to DNA. Therefore, we first attempted to establish a comparative characterization of the energetics and dynamics of one initial representative model from each group.

Following 2 μs of CG molecular dynamics on these systems, it was observed that all three systems displayed a decrease in alpha helix and extended beta sheet content comparing to the relaxed capsid structure, with the most pronounced reduction occurring in the disordered system (**Figure 3A**). An analysis of the final frame of the simulations indicated that the capsid exhibited changes in morphology and internal arrangements when comparing the three systems (**Figure 3B**). These differences suggest that varying genome geometries produce differences in capsid topology. To address this initial observation and enhance conformational diversity sampling, we constructed multiple models within each of the three topological categories.

**Figure 3.**
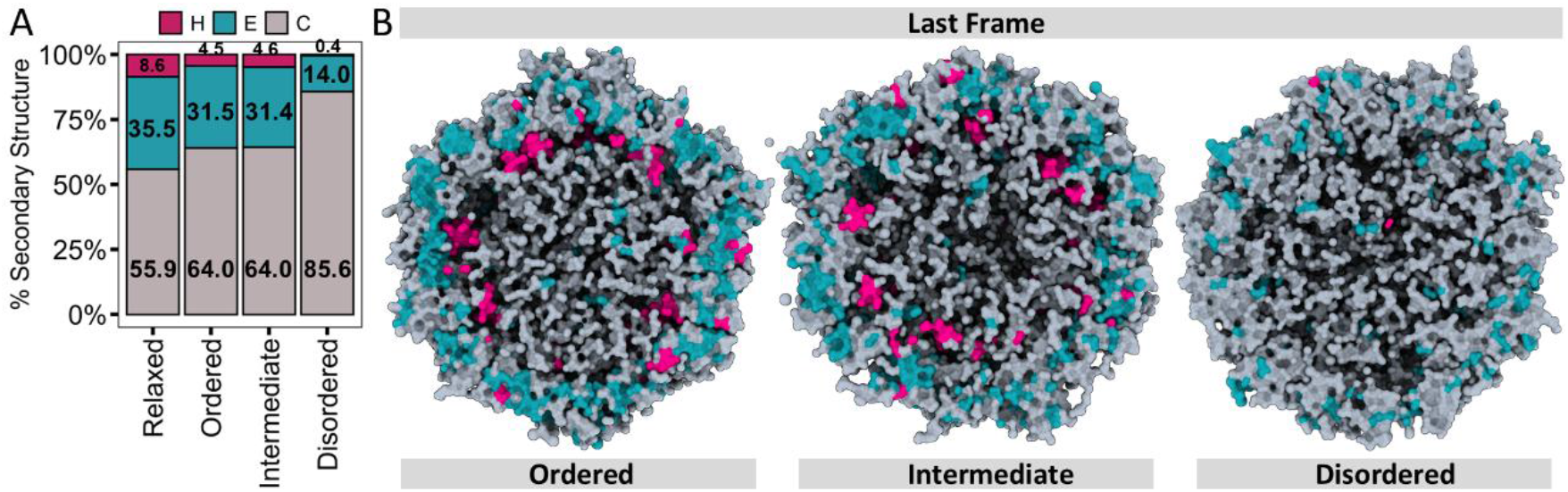
Capsid topology after CG molecular dynamics simulation. A) The secondary structure composition during the CG SIRAH simulations was assessed across the 2 μs duration for the Ordered, Intermediate, and Disordered systems. The secondary structure content of the relaxed model was also computed for comparison purposes. Coloring followed SIRAH secondary structure classification where alpha helices (H) are pink, extended beta sheets (E) are blue, and coils (C) are gray. B) Cut-view images of the capsid structure at the end of the simulations. Surface colors as in A.

To ensure different conditions, three ordered models were generated by manually constructing a Hamiltonian path where all Cap pentagons were visited. In addition, we employed different protocols for compacting the DNA inside the capsid at the SPQR level. This choice allowed us to obtain different geometries and different packaging densities, giving rise to three ordered models, O1, O2 and O3. Model O1 was compacted to a radius of gyration to 4.801 nm with a constant of 10000 *ϵ*_*s*_/Å^2^, along a series of 5000 short simulations of 20000 MC sweeps, which minimized the energy and simultaneously removed the clashes, acting only on the clashed fragments. Model O2 employed the procedure described in the Methods section, while model O3 was constructed from model O2 but additionally expanding the radius of gyration through another simulation of 100000 steps, with an equilibrium radius of gyration of 4.32 nm and a constant of 10000 *ϵ*_*s*_/Å^2^.

Intermediate model I1 was created manually, imposing only 6 complete pentagons along its path, while I2 was obtained with the MC protocol, containing only one full pentagon. Additionally, four new disordered models were using the MC protocol described in the Methods section. All constructed genomic topologies are depicted in **Supplementary Figure S2** and will henceforth be represented in varying shades of blue, orange, and red, corresponding to Ordered, Intermediately Ordered, and Disordered topologies, respectively.

As it will be shown below, all models resulted in similar sizes well within experimental determinations. Hence, we will not go through detailed descriptions of the interactions in each model, but rather provide a comparative characterization of the different topological groups.

### Disordered topologies increase structural distortions without compromising virion sizes

CG molecular dynamics simulations showed that all systems slowly stabilized after approximately 1 μs. While all systems showed similar kinetics, the capsids of virions with disordered models exhibit larger structural deviations from their starting points, as evidenced by the clearly separated RMSD traces shown in **Figure 4A**. However, the same behavior is not retrieved when considering the mobility of the genomes. Although Disordered models showed higher RMSD values, there are at least two remarkable differences (**Figure 4B**). First, the convergence is much faster for the DNA filaments, suggesting a very fast rearrangement of the genomes takes place soon after the beginning of the simulations. Second, the final RMSD values of the genomes roughly double those of the protein capsid. This suggests that, despite the genome topology being determinant for the capsid’s stability, the global dynamics of both components are somehow decoupled. This can be related to the lack of secondary structure elements in the DNA and the N-termini of the Cap proteins, while the more external protein shell is highly structured and compact (23).

**Figure 4.**
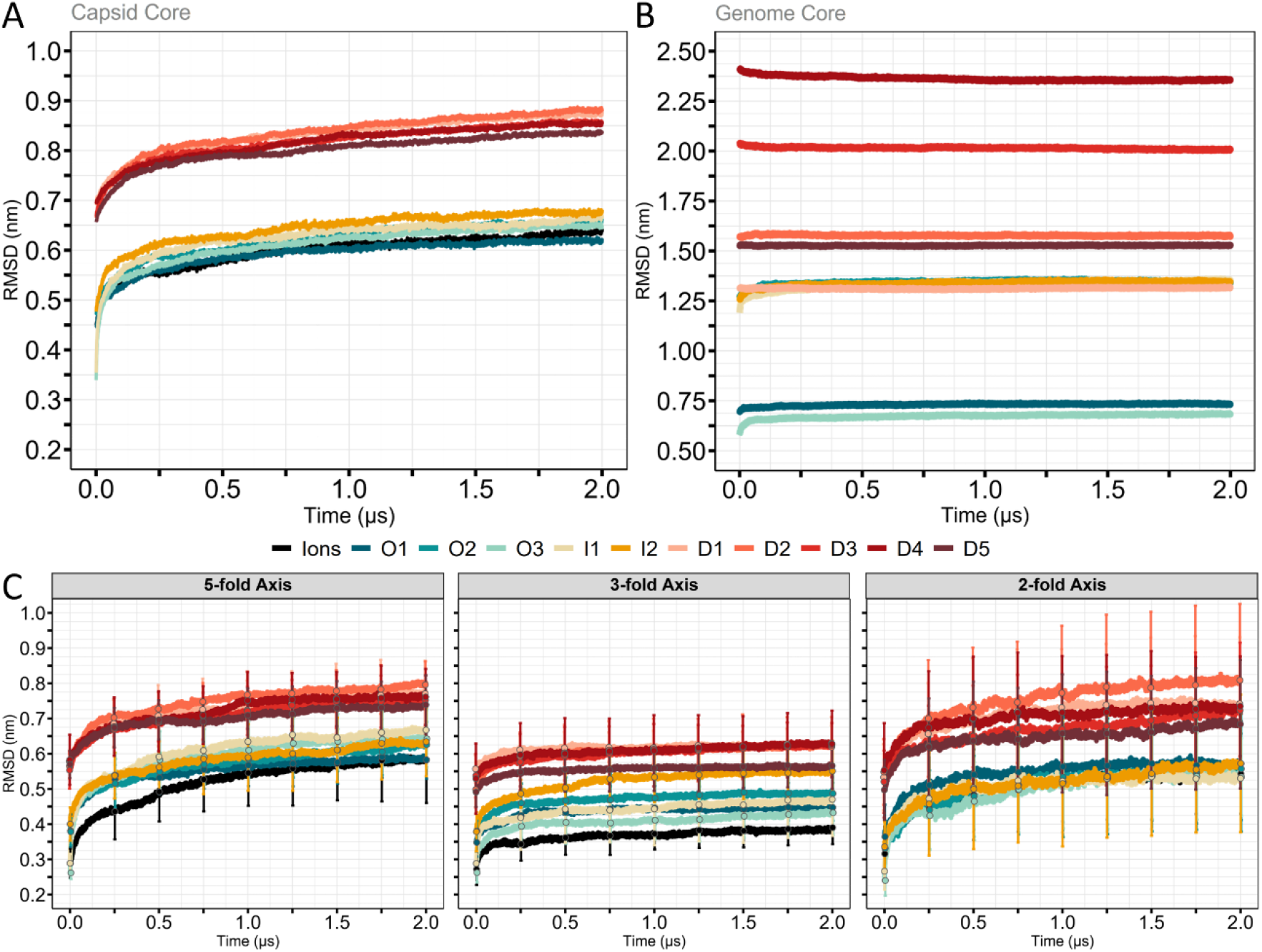
Topological diversity in PVC2d virions results in different dynamics with similar sizes. A) RMSD traces on the protein capsid for the different models. B) RMSD traces on the Phosphate beads of the genome for the different models. C) RMSD calculated on the beads corresponding to the neighborhood of the symmetry axis. From left to right, data is averaged over twelve 5-fold axes, twenty 3-fold axes, and thirty 2-fold axes. The standard deviations for each trace are shown as error bars at only 10 time points for visual clarity.

Aimed to set a base line dissect the DNA topology, we also constructed a model without DNA but with 1769 Chlorine ions distributed uniformly inside the capsid and the same simulation protocol was repeated. In this way, we could separate the electrostatic stabilization from the effects of the topology. The conformational behavior of this model is indicated by a black line in **Figure 4**. Remarkably, the RMSD traces of this model are nearly indistinguishable from those of models O1, O2, and O3, supporting the idea that the Ordered topologies are optimized in terms of structural stability and their cross talk with the proteinaceous parts is well balanced.

Since the packaging signals are located in the close neighborhood of the 3-fold symmetry axis, and previous simulations of the PCV2d virus-like particle containing only the tetranucleotides bound to the packaging signals revealed a stabilizing effect in that region (52), we wonder if the presence of the entire genome would exert a similar effect. Analyzing the structural distortions in the vicinity of the symmetry axis revealed a differential behavior in this case as well. The structural deviations show diverse patterns around the symmetry axis, with the 3-fold displaying a faster convergence and less pronounced differences between the ordered, intermediately ordered, and disordered topologies (**Figure 4C**). This would be consistent with a DNA-mediated stabilization of the vicinity of the 3-fold axis and a relatively compact structural motif observed in that region of the capsid. Additionally, we noticed that the neighborhood of the 2-fold axis exhibits much higher variability, as evidenced by the larger standard deviations observed (**Figure 4C**). Also in these descriptors, the cations-only system is well comparable to those of the Ordered topologies.

In contrast to deviations from starting configurations, the diameters estimated along MD trajectories show no evident correlation with the topology and all simulations sample very similar values, with differences below 0.4 nm among themselves and very close to the CryoEM value (**Figure 5A**). These small differences are very well within a recent diameter determination of 19.4 nm +/− 3 nm made by our group using Atomic Force Microscopy (52).

**Figure 5.**
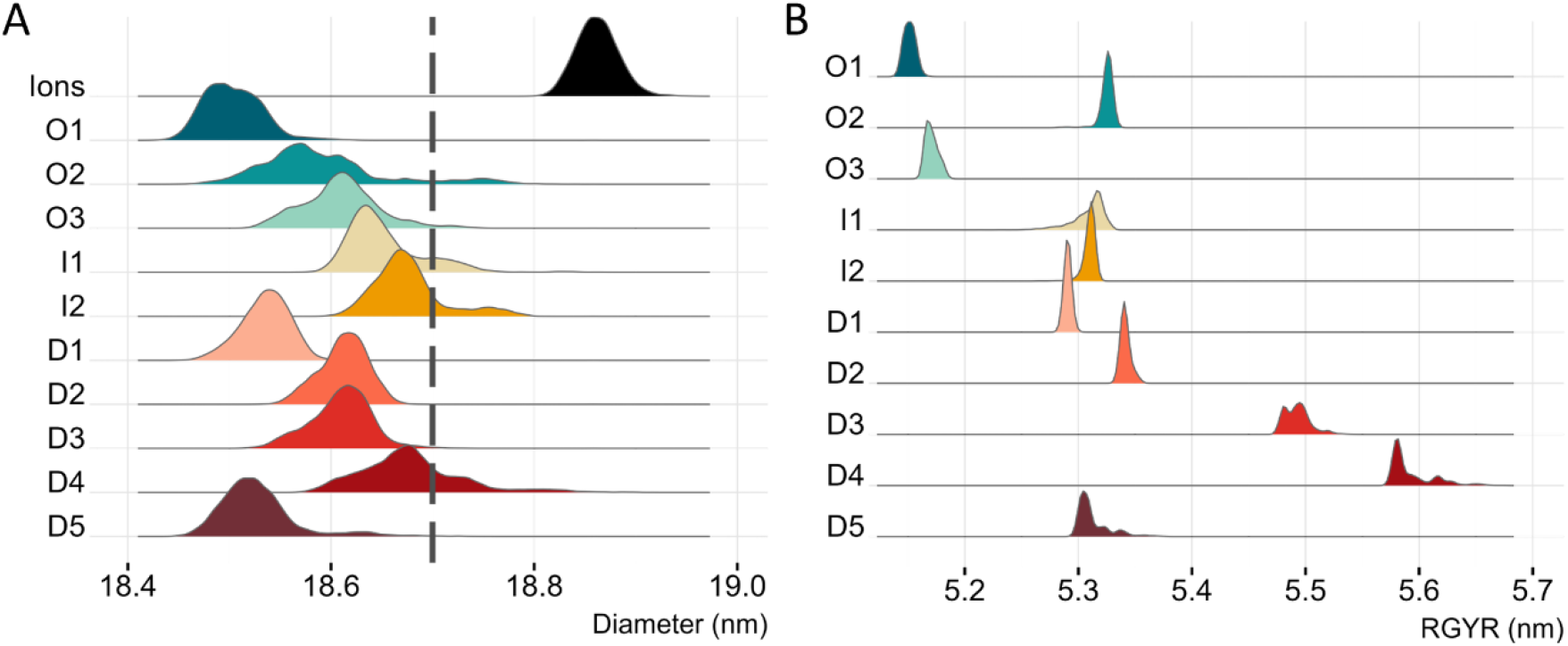
Average sizes of protein and DNA components. A) Distribution of the external capsid diameters measured along the MD trajectories. B) Distribution of the radius of gyration of the genomes along the MD trajectories.

Unfortunately, structural data about the global size of the ssDNA genome folded inside the capsid is not available. However, it is possible to achieve a comparative gauge by comparing the gyration radii on the different systems simulated. As it can be seen from **Figure 5B**, all gyration radii display a very narrow distribution in comparison to that seen for the protein capsid (**Figure 5A**) and there is no clear correlation between the average sizes of the proteic and nucleic components of the virion. Again, this suggests the idea of a decoupled dynamics between the internal and external parts of the virion. Alternatively, it is possible that the averaging intrinsic in the gross determinants reported in **Figures 4 and 5**. To analyze this possibility, we turned our attention to the details of protein-DNA interactions.

### Ordered genomes keep native contacts

The rapid loss of symmetry during the dynamics is not surprising, given the temperature effects. Moreover, the conformations of the modeled genomes are intrinsically asymmetric. However, the clear separation of structural deviations and the uncorrelated conservation of virion sizes were unexpected and prompted us to explore the preservation of local interactions. First, we calculated the percentage of native protein-protein contacts maintained during the simulations in the vicinity of the symmetry axis, using the same residue groups used in the RMSD calculations in **Figure 4**. Consistent with our previous observations, different topologies yield distinct behaviors. Disordered topologies experience the highest loss of native contacts (**Figure 6A**). In all cases, the differences between the traces indicate a lower native contact conservation by above 10% in the Disordered virions, when compared against other topological arrangements in the three symmetry axes. Notably, the largest deviations occur in native contacts around the 2-fold axes, which can be attributed to the low secondary-structure content in this region.

**Figure 6.**
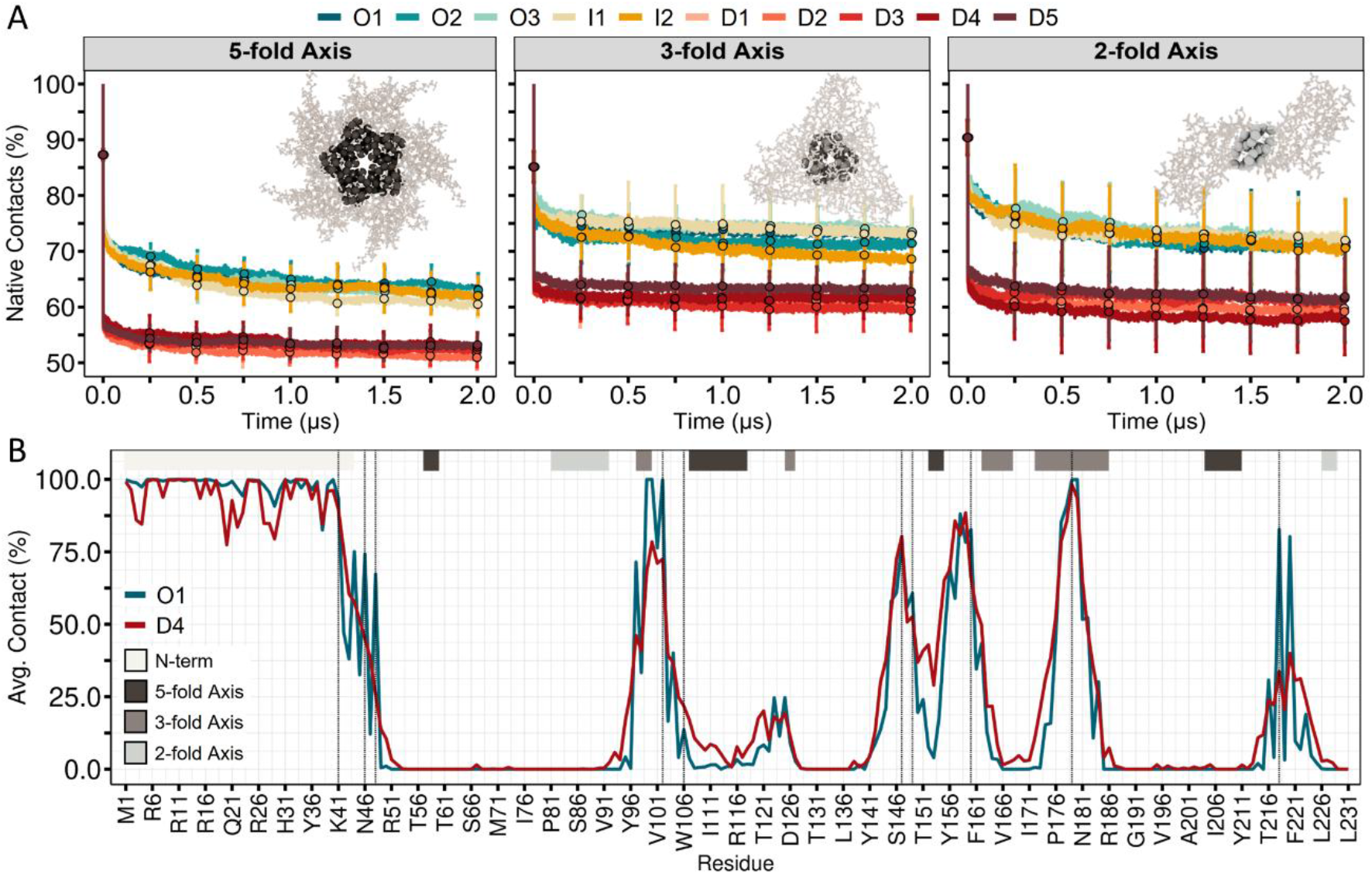
Genome ordering influences protein-protein and DNA-protein contacts. A) Percentage of native contacts conservation during the simulations in the neighborhood of the symmetry axis. The standard deviations for each trace are shown as error bars at only 10 time points for visual clarity. B) Average DNA-protein contacts. A contact is counted if a nucleotide is found within 0.6 nm of any amino acid, and averaged over the MD trajectory and the 60 capsomers. The regions corresponding to the symmetry axis in the Cap protein are indicated at the top of the panel. Vertical dashed lines correspond to experimental contact measured in the CryoEM structure. Only Models O1 and D4 are shown for visual clarity, but the same features are valid for the ten models simulated, according to their ordering (see **Supplementary Figure S3**).

Subsequently, we turned our attention to the DNA-protein interactions. We calculated DNA-protein contacts, averaging over the 60 copies of the capsid protein to obtain a percentage of contacts between nucleotides and amino acids.

As shown in **Figure 6B**, the N-terminal regions, protruding into the virion’s center, are closely surrounded by nucleic acids with a very high percentage of DNA contacts during the dynamics. After Lysine 41, there is an alternation of peaks and basins, indicating regions of the protein alternating between DNA contacts, protein-protein contacts, or amino acids exposed to the exterior. The good correspondence between the peaks and the dashed lines, corresponding to experimental contacts, suggests faithful conservation of the global interactions. However, comparing ordered vs. disordered genomes reveals that DNA-protein interactions are better maintained in the Ordered topologies (only two are shown for visual clarity, all topologies are shown in **Supplementary Figure S3**). Notably, Ordered models not only feature better average conservation of experimental contacts, but also show the N-terminal region consistently higher and more homogeneous values. This may suggest that genome visiting all packaging signals in an orderly fashion may result in a better stabilization of DNA-protein contacts even in intrinsically disordered regions located several nm away from the packaging signal.

### Disordered genomes are less stable and exhibit lower melting temperatures

Finally, we sought to analyze whether the differential structural and dynamical behavior observed for different genome arrangements may translate into dissimilar stabilities.

To gain insights into the thermal stability of the different virion topologies, we first calculated the potential energy of the entire systems, including DNA-protein-solvent interactions. As seen in **Figure 7A**, all systems are stable (as indicated by negative energy) at 27 °C. Also in this case, all energy traces present a monotonic and similar increase, indicating an analogous stabilization in all cases. However, virions with Organized and Intermediate genomes are well separated from Disordered ones, establishing a direct link between the topological organization and the stability of the resulting viral particles. Aimed to dissect the solvent contribution, we calculated only the DNA-protein contribution. As this quantity may be more sensitive to conformational changes during relaxation, we decided to average these energies over the entire simulations. Stinkingly, the internal energy of some of the Disordered virions is marginally stable or even positive (**Figure 7B**). Therefore, for some of these genome-Disordered viruses, solvation plays a crucial role that can even counteract internal repulsions.

**Figure 7.**
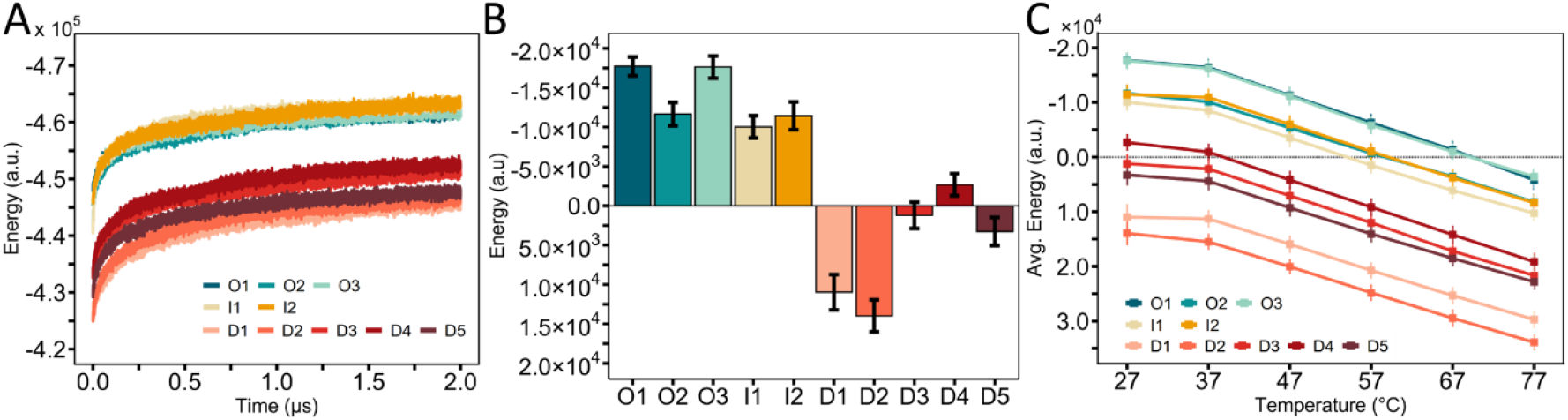
Genome ordering and virion energetics. A) Total energy of the entire systems throughout the simulations. B) DNA-protein interaction averaged over each of the simulations. C) DNA-protein averaged over the heating simulations. The datapoints correspond to the energy averaged over the corresponding 10-degree interval. As the temperature rises, solute energy diminishes in all systems. The melting point is the temperature at which the energy crosses zero.

The results of the energetic analysis show that genetically identical but topologically distinct virions may exhibit significantly different stabilities, which should translate into distinct melting points. To explore that possibility, we used the final conformation from each simulation and coupled each system to progressively higher temperatures in steps of 10 °C. During each of these heating steps, we calculated the DNA-protein energy as the average over that simulation period. In line with our speculation, the ordering in the encapsidated genomes not only separated the energy values but also yielded nearly parallel, monotonically decreasing energy values with increasing temperature (**Figure 7C**). The melting temperature corresponds to the point at which these lines cross the zero-energy level.

At this point, it might be relevant to recall that PCV2 is highly thermostable with inactivation temperatures of nearly 75 °C (61). So, it is remarkable to find that the melting point featured by our Ordered and Intermediately ordered virions is in the range of the inactivation temperature reported for intact virions.

## Discussion and Conclusions

We have introduced a systematic, multiscale approach to explore and control the global arrangement of the genome, starting from limited knowledge about the DNA-capsid contacts. The models can be massively produced and easily refined up to pseudoatomistic resolution through a progressive incorporation of chemical details. The lattice representation provides a better understanding of the global features which, as we showed here, are relevant for the stability and structural properties of the viruses, and can be easily extended to other topologies, to include, for example, secondary structure elements such as hairpins and junctions. Moreover, this sampling method and the backmapping procedure can be straightforwardly applied to other viral structures where the structure of fragments of the genome close to the capsid are known (62).

From a virological perspective, our results provide new insights into viral variability and stabilization. The theory of viral quasispecies was originally developed to describe populations of rapidly replicating viruses as dynamic distributions of closely related genetic variants rather than as single, well-defined genotypes (63). In this framework, the capacity of a virus to replicate (namely, its fitness) is not an intrinsic property of an individual sequence, but an emergent property of the entire population, shaped by mutation, selection, and replication dynamics. Our results suggest a possible extension of the quasispecies concept beyond genetic variation to encompass other sources of heterogeneity that affect viral performance. We showed that even in the absence of mutations, structurally identical genomes may adopt energetically viable multiple conformations when confined within a capsid, giving rise to an ensemble of particles with distinct physical and functional properties. These alternative genome organizations may translate into differences in thermal stability, packaging efficiency, or uncoating readiness, thereby contributing unequally to viral fitness. From this perspective, the conformational variability of the packaged genome can be viewed as a quasispecies-like distribution in structural space, where selection acts on particle-level phenotypes rather than on nucleotide-sequence differences.

Recently, a thorough study followed the spontaneous mutations that arose in a cohort of piglets infected with PCV2 (64). Animals were divided into three groups according to whether they developed null, mild, or severe symptoms. Measuring the number of mutations after the first, second, and third weeks post-infection resulted in a significant drop in the number of spontaneous mutations to virtually zero (see Figure 2 in Ref. (64)).

Our results suggest that even if a strong selective pressure exerted by the immune system limits the mutational space, compact viruses, such as Circoviruses, can still add a new dimension in variability, which favors the success across replication cycles by creating an ensemble of conformations with different functional biases as uncoating efficiency or disassembly under temperature variations.

Under a thermodynamic perspective, the population of genome-ordered/disordered virions would behave as an ensemble of “conformational quasispecies” that could provide a physics-based source of variability when biological options are exhausted. Furthermore, the structure of the topologies studied here may also correlate with the efficiency of the assembly pathway, distinguishing cases in which local pentamer assembly precludes global viral assembly, as pointed out by studies in similar (36).

## Conflict of interest

The authors declare no competing financial interest.

## Funding

This work was funded by FOCEM (MERCOSUR Structural Convergence Fund), COF 03/11. Lucianna H. Silva Santos and Sergio Pantano are members of the Uruguayan SNI. Financial support by ANII through the Maria Viñas Fund FMV_1_2023_1_175988 is greatly acknowledged. Simón Poblete was funded by ANID Fondecyt Regular project No. 1231071 and Centro Ciencia & Vida, FB210008, Financiamiento Basal para Centros Científicos y Tecnológicos de Excelencia de ANID (https://anid.cl).

## Notes

### Competing Interest Statement

The authors have declared no competing interest.

