## Supplementary material text and figures. for "Topology-aware multiscale modeling of viral genomes reveals stability determinants in circoviruses"

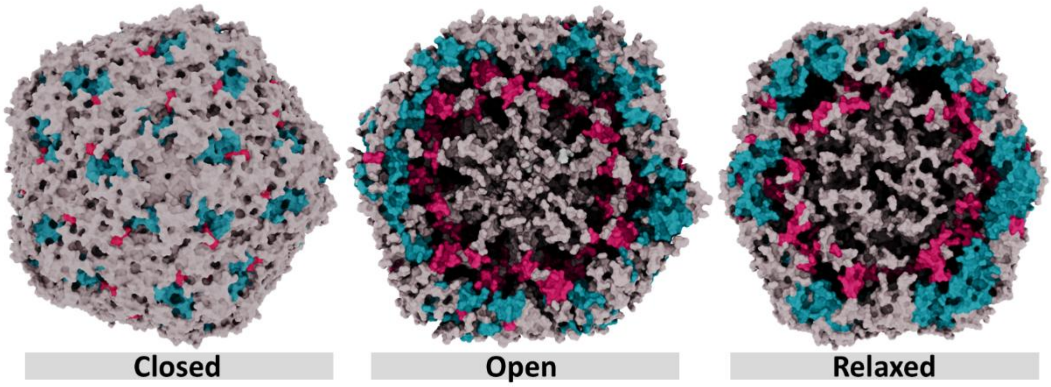


**Figure S1.** The surface representation of the modeled PCV2 capsid is color-coded according to secondary structure: pink for alpha helices, blue for extended beta sheets, and gray for coils. The closed and open representations depict the model with the predicted Cap protein, whereas the relaxed representation illustrates this capsid following minimization and relaxing processes via molecular dynamics (MD) simulations.

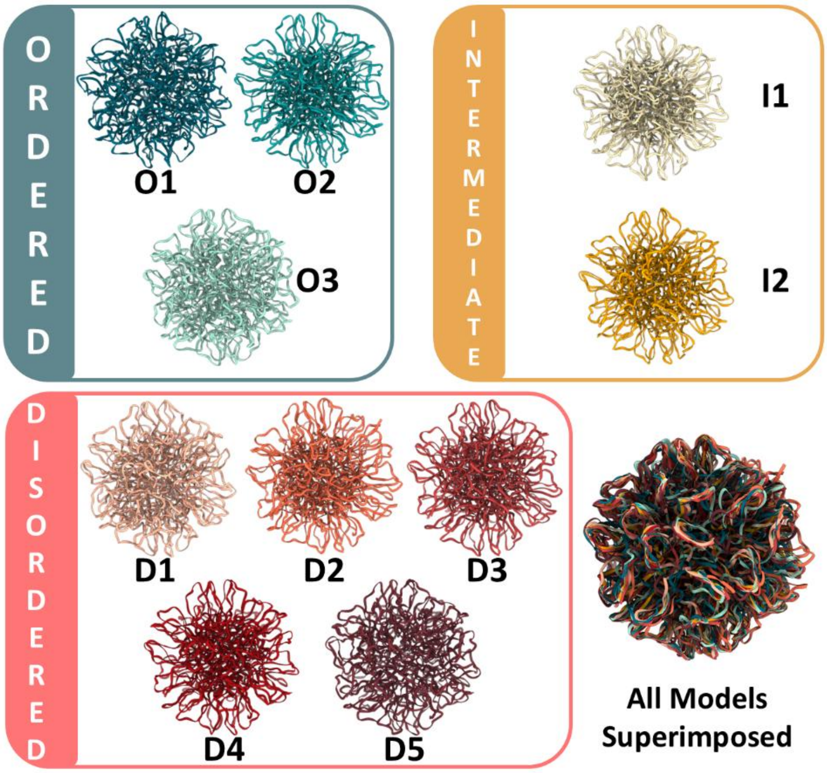


**Figure S2.** Pictorial view of the different genome topologies generated. Different tones of green, orange, and red colors correspond to Ordered, Intermediately ordered, and Disordered topologies, respectively. The same color coding is used throughout the main manuscript.

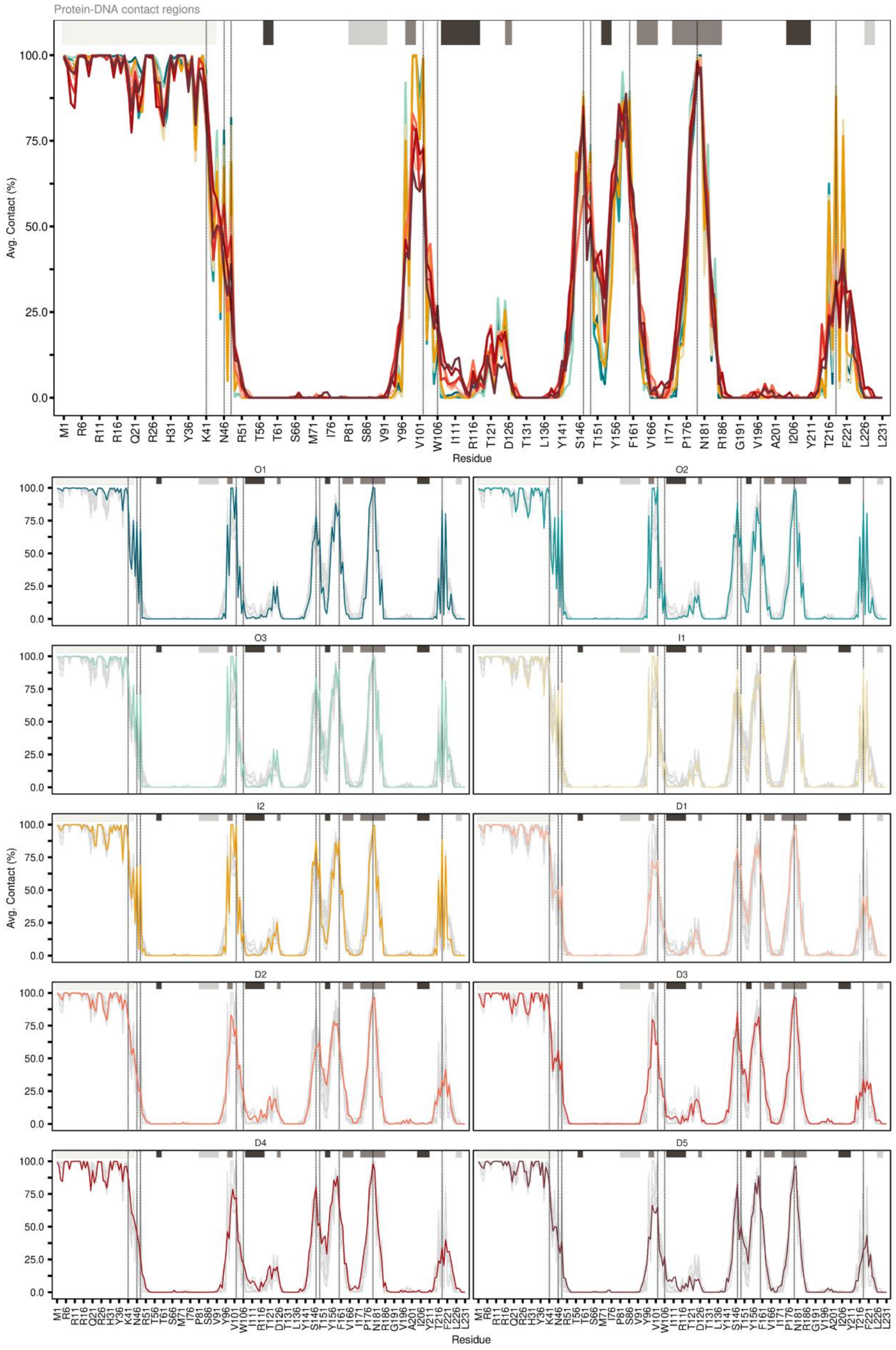


**Figure S3.** Average DNA-protein contacts versus the amino acid number. The top panel shows the contacts for all models. Note that disordered models (redish colors) display a lower conservation. The gray segments on the top indicate the secondary structure of the Cap protein. The dashed lines indicate amino acids forming DNA-protein contacts in the cryo-EM structure. The same information or each of the models simulated is shown separately in the bottom panels.
